# Obesity alters *Ace2* and *Tmprss2* expression in lung, trachea, and esophagus in a sex-dependent manner: Implications for COVID-19

**DOI:** 10.1101/2020.10.13.337907

**Authors:** Dylan C. Sarver, G. William Wong

## Abstract

Obesity is a major risk factor for SARS-CoV-2 infection and COVID-19 severity. The underlying basis of this association is likely complex in nature. The host-cell receptor angiotensin converting enzyme 2 (ACE2) and the type II transmembrane serine protease (TMPRSS2) are important for viral cell entry. It is unclear whether obesity alters expression of *Ace2* and *Tmprss2* in the lower respiratory tract. Here, we show that: 1) *Ace2* expression is elevated in the lung and trachea of diet-induced obese male mice and reduced in the esophagus of obese female mice relative to lean controls; 2) *Tmprss2* expression is increased in the trachea of obese male mice but reduced in the lung and elevated in the trachea of obese female mice relative to lean controls; 3) in chow-fed lean mice, females have higher expression of *Ace2* in the lung and esophagus as well as higher *Tmprss2* expression in the lung but lower expression in the trachea compared to males; and 4) in diet-induced obese mice, males have higher expression of *Ace2* in the trachea and higher expression of *Tmprss2* in the lung compared to females, whereas females have higher expression of *Tmprss2* in the trachea relative to males. Our data indicate diet- and sex-dependent modulation of *Ace2* and *Tmprss2* expression in the lower respiratory tract and esophagus. Given the high prevalence of obesity worldwide and a sex-biased mortality rate, we discuss the implications and relevance of our results for COVID-19.

## INTRODUCTION

Since the worldwide spread of COVID-19 caused by SARS-CoV-2 [1–4], epidemiological studies have highlighted obesity as a major risk factor for development of COVID-19, disease severity, and death [5–10]. This association is not merely due to the high prevalence of obesity among world populations. A meta-analysis of 75 published studies from more than 10 countries in Asia, Europe, and North and South America confirmed a link between obesity and COVID-19 [9]. However, how obesity influences and contributes to the pathophysiology of COVID-19 is hotly debated— multiple mechanisms have been suggested, ranging from obesity-linked alteration in lung function, vascular health, systemic low-grade inflammation, altered immune response, and viral-bacterial interactions [11–17].

SARS-CoV-2 uses the host cell transmembrane carboxypeptidase ACE2 as a receptor to enter cells [18–20]. This process relies on the type II transmembrane serine protease TMPRSS2 to cleave and prime the viral spike protein for cell entry [18]. Expression of both ACE2 and TMPRSS2 in target cells is thought to be important for viral infection. Data from single-cell RNA sequencing reveal expression of *ACE2* and *TMPRSS2* in multiple cell types within the heart, lung, trachea, esophagus, kidney, gastrointestinal tract, and liver [21–24]. Since SARS-CoV-2 is a respiratory virus that causes significant lung pathology [25] and is spread through person-to-person contact [26] or aerosolized nasal droplets [27, 28], viral entry through the respiratory tract is thought to be the major route of infection.

In the present study, we hypothesized that obesity alters expression of *ACE2* and *TMPRSS2* in the lower respiratory tract and that these changes could potentially contribute to greater risk of developing COVID-19. To test this, we evaluated expression of *Ace2* and *Tmprss2* in the trachea, lung, and esophagus of chow-fed lean mice and obese mice of similar age chronically fed a high-fat diet (HFD). We observed both diet- and sex-dependent modulation of *Ace2* and *Tmprss2* expression in the lower respiratory tract (trachea and lung) and esophagus. Given the high prevalence of obesity worldwide [29] and sex biases in COVID-19–related death [30], this study has relevance and implications for COVID-19.

## MATERIALS AND METHODS

### Mice

C57BL/6J male and female mice were housed in polycarbonate cages on a 12-h light-dark photocycle with ad libitum access to water and food, with no more than five adult mice per cage. Mice were fed either standard laboratory chow (Tekland 2018SX, Envigo, Indianapolis, IN) or a HFD (60% kcal derived from fat; D12492, Research Diets, New Brunswick, NJ) beginning at 6 weeks of age. At termination of the study, food was removed for 2 h before euthanasia. All animal protocols were approved by the Institutional Animal Care and Use Committee of The Johns Hopkins University School of Medicine (Protocol # MO16M431).

### Body composition analysis

Body composition analyses for total fat and lean mass were determined using a quantitative magnetic resonance instrument (Echo-MRI-100, Echo Medical Systems, Waco, TX) in the Mouse Phenotyping Core facility at Johns Hopkins University School of Medicine.

### Tissue collection

Whole lung, trachea, and esophagus were immediately harvested from euthanized mice, flash-frozen in liquid nitrogen, and stored at –80°C until analysis.

### Quantitative real-time PCR

Total RNA was isolated from lung, trachea, and esophagus using Trizol reagent (Life Technologies, Frederick, MD), reverse transcribed using the iScript cDNA synthesis kit (Bio-Rad, Hercules, CA), and subjected to quantitative real-time PCR analyses using SsOAdvanced Universal SYBR Green Supermix (Bio-Rad) on a CFX Connect Real-Time System (Bio-Rad). A total of 50 ng (lung samples) or 100 ng (trachea and esophagus samples) of reverse-transcribed cDNA were used in each real-time PCR reaction. Data were normalized to the average of two house-keeping genes (β-actin and cyclophilin A) and expressed as relative mRNA levels using the ΔCt method [31]. Real-time PCR primers used in this study were: *Ace2* forward, 5’-TCCAGACTCCGATCA TCAAGC-3’ and reverse, 5’-GCTCATGGTGTTCAGAATTGTGT-3’; *Tmprss2* forward, 5’-CAGTCTGAGCACATCTGTCCT-3’ and reverse, 5’-CTCGGAGCATACTGAGGCA-3’; *β-actin* forward, 5’-AGTGTGACGTTGACATCCGTA-3’ and reverse, 5’-GCCAGAGCAGTAATCTCC TTCT-3’; *cyclophilin A* forward, 5’-GAGCTGTTTGCAGACAAAGTTC-3’ and reverse, 5’-CCC TGGCACATGAATCCTGG-3’.

### Statistical analysis

All data are presented as mean ± SEM. Single-variable comparisons between two groups of data were performed using two-tailed Student’s *t-*test with 95% confidence intervals (i.e. male chow vs male HFD, in which gene expression is measured and diet is the variable; and male chow vs female chow, in which gene expression is measured and sex is the variable). Comparisons were not made across two variables, for example, male chow vs female HFD. Prism 8 (GraphPad Software, La Jolla, CA, USA) was used for statistical analyses, and differences were considered to be statistically significant at *P* < 0.05.

## RESULTS

Age, body weight, total fat and lean mass, and lung weight of male and female mice fed control chow or a HFD are indicated in Table 1. As expected, five months of chronic high-fat feeding resulted in significant weight gains and fat mass accrual in both male and female mice relative to control chow-fed mice.

**Table 1.** Age, sex, weight, total fat and lean mass, and lung weight of lean and obese mice.

|  | Male |  | *P* |  | Female |  | *P* |
| --- | --- | --- | --- | --- | --- | --- | --- |
| *N* | 9 | 8 |  |  | 10 | 10 |  |
| Diet | Chow | High-fat diet |  |  | Chow | High-fat diet |  |
| Age (months) | 6 | 6.5 |  |  | 6 | 6.5 |  |
| Weight (g),<br>mean ± SEM | 29.0 ± 0.45 | 45.2 ± 1.49 | <0.0001 | | 21.9 ± 0.76 | 36.5 ± 1.50 | <0.0001 |
| Total fat mass (g),<br>mean ± SEM | 4.80 ± 0.46 | 18.6 ± 0.94 | <0.0001 | | 2.60 ± 0.23 | 12.6 ± 0.98 | <0.0001 |
| Total lean mass (g),<br>mean ± SEM | 21.1 ± 0.41 | 20.8 ± 0.37 | 0.600 | | 17.2 ± 0.44 | 17.1 ± 0.34 | 0.907 |
| Lung weight (g),<br>mean ± SEM | 0.148 ± 0.002 | 0.159 ± 0.004 | 0.017 | | 0.140 ± 0.005 | 0.156 ± 0.003 | 0.008 |

### Obesity alters expression of *Ace2* and *Tmprss2* in lung, trachea, and esophagus

Quantitative real-time PCR showed that in obese male mice fed an HFD, expression of *Ace2* was significantly elevated in the lung and trachea relative to chow-fed lean male mice (Fig. 1A,C). In esophagus, *Ace2* expression did not significantly differ between obese and lean male mice (Fig. 1E). In obese female mice, however, expression of *Ace2* in the esophagus was significantly reduced relative to lean female animals (Fig. 1E). In contrast to *Ace2* expression, *Tmprss2* expression was significantly reduced in the trachea of obese male mice relative to lean controls (Fig. 1D). In obese female mice, *Tmprss2* expression was significantly reduced in the lung but elevated in the trachea compared to female lean controls (Fig. 1B,D).

**Figure 1.**
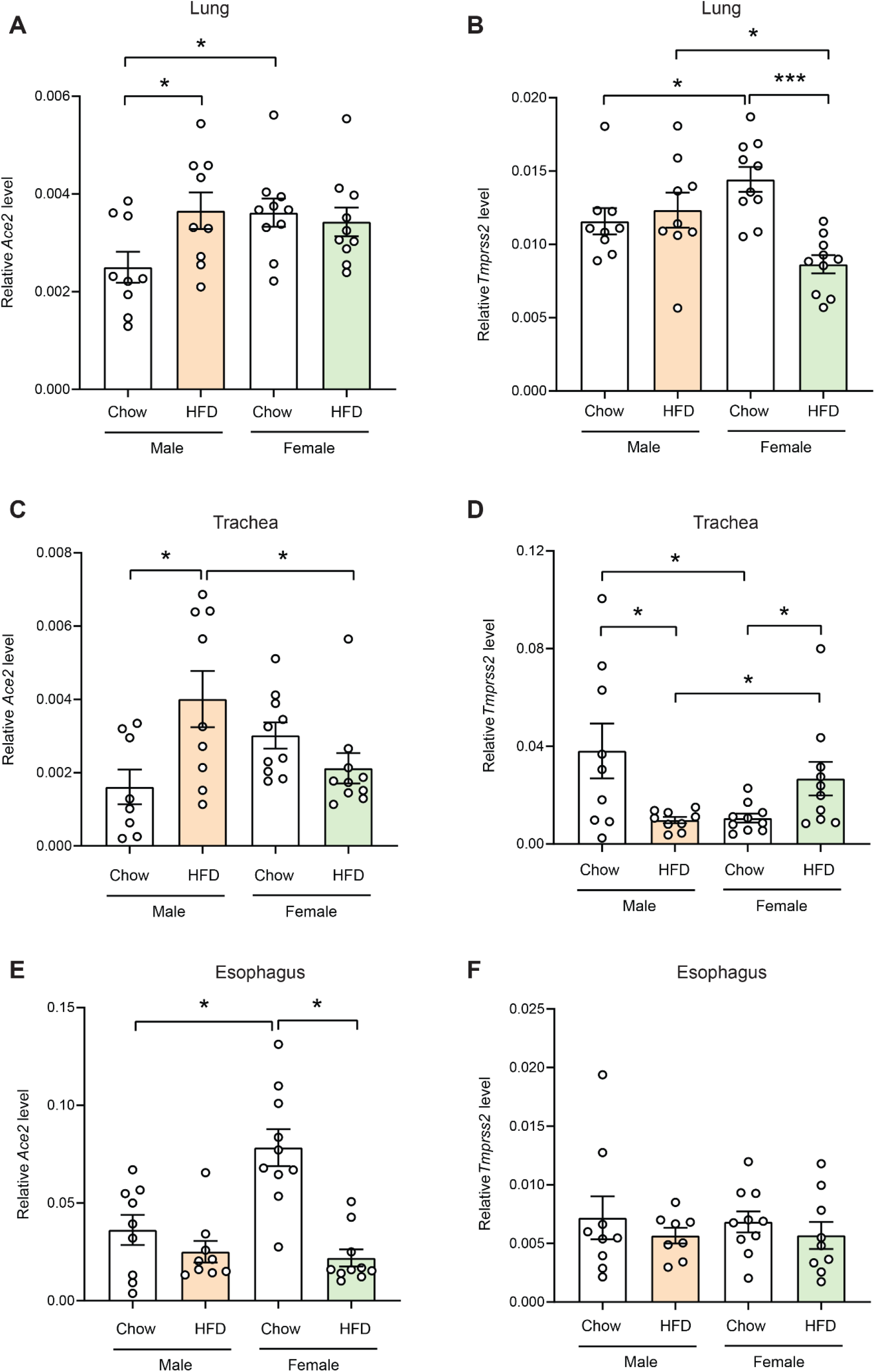
Expression of *Ace2* and *Tmprss2* in lean and obese mice. Real-time PCR analysis of *Ace2* and *Tmprss2* transcript levels in lung (A, B), trachea (C, D), and esophagus (E, F) of lean male (*n* = 9), obese male (*n* = 8), lean female (*n* = 10), and obese female (*n* = 10) mice. All expression data were normalized to the average of two house-keeping genes (*β-actin* and *cyclophilin A*). All data are expressed as mean ± SEM. * *P* < 0.05; *** *P* < 0.001

### Sex differences in *Ace2* and *Tmprss2* expression in chow-fed lean mice

In chow-fed lean mice, females had significantly higher expression of *Ace2* in the lung and esophagus than males (Fig. 1A,E). Although expression of *Ace2* in trachea also trended higher in lean females compared to lean males, the differences were not significant (Fig. 1C). Interestingly, chow-fed female mice had significantly higher *Tmprss2* expression in the lung but significantly lower expression in the trachea compared to chow-fed male mice (Fig. 1B,D). Unlike in the lower respiratory tract, no sex differences were noted in expression of *Tmprss2* in the esophagus of chow-fed animals (Fig. 1F).

### Sex differences in *Ace2* and *Tmprss2* expression in HFD-fed obese mice

When comparing obese male and female mice, males had significantly higher expression of *Ace2* in the trachea and *Tmprss2* in the lung than females (Fig. 1B,C). Obese female mice, however, had significantly higher expression of *Tmprss2* in the trachea relative to obese male mice (Fig. 1D).

## DISCUSSION

Our data indicate sexually dimorphic expression patterns of *Ace2* and *Tmprss2* in both lean and diet-induced obese mice. Although it is not known at present whether expression of *ACE2* and *TMPRSS2* in the lung, trachea, and esophagus of humans also exhibit similar patterns of sexual dimorphism, our data have relevance and implications for COVID-19.

Initial viral loads are thought to impact the severity of COVID-19. Indeed, patients with more severe COVID-19 have higher viral loads in the respiratory tract (throat, bronchoalveolar lavage fluid, or sputum) and longer viral persistence than those who experience milder disease [32–34]. Not surprisingly, expression levels of the viral host receptor ACE2 and cell entry-associated molecules (e.g., TMPRSS2) are thought to be important and relevant factors influencing viral loads and infection. Although elevated expression of ACE2 in the upper airway (nasal and olfactory epithelium) may be linked to anosmia in patients with COVID-19 [35, 36], its expression in the lung appears to plays a crucial role in SARS-CoV-induced lung injury in animal models [37].

Of the comorbidities associated with COVID-19, obesity consistently ranks among the highest risk factors influencing COVID-19 diagnosis (>46% higher), hospitalization (113% higher), ICU admission (74% higher), and mortality (48% higher) based on a meta-analysis of 109,367 individuals from 75 published studies from more than 10 countries [9]. Obesity also appears to shift COVID-19 severity to younger ages [8]. Epidemiological data further suggest that men have ~2.5% greater mortality due to COVID-19 than women [30].

How do sex and obesity influence the pathobiology of COVID-19? Many host factors, both genetics and behavioral, are likely at play, with different degrees of importance and contribution to overall disease severity and viral persistence [11–17]. The present study sought to address one basic question—does obesity alter expression of *Ace2* and *Tmprss2* in the lower respiratory tract and esophagus of male and female mice? Our data show that obese male mice indeed had significantly higher *Ace2* expression in the lung than lean controls. Although obese female mice had comparable *Ace2* levels in the lung as female lean controls or obese male mice, *Tmprss2* transcript levels in the lung were significantly reduced relative to lean controls and obese male mice.

The trachea connects the upper airway (nose and throat) to the lung and thus represents an additional high virus-to-tissue contact zone. In this organ, *Ace2* expression was also significantly higher in obese male mice relative to lean male controls or to obese female mice. Together, these observations may potentially account, at least in part, for the association of obesity with SARS-CoV-2 infection and COVID-19 severity, as well as the male-biased mortality rate. Interestingly, expression of *Tmprss2* in trachea was significantly lower in obese male mice relative to lean male controls and obese female mice. It has been suggested that other cellular proteases (e.g., furin and cathepsin B/L) may potentially be used by SARS-CoV-2 to enter *Tmprss2*-negative cells [18, 38]. Thus, elevated *Ace2* viral receptor expression in the trachea of obese male mice, despite lower expression of *Tmprss2*, may still favor viral entry if other cellular proteases can substitute for Tmprss2.

Some limitations of the present study are noted, however. (1) Only *Ace2* and *Tmprss2* transcript levels were assessed, and we presumed that ACE2 and TMPRSS2 protein abundance correlated with their mRNA levels. (2) Quantifications of *Ace2* and *Tmprss2* transcript levels were performed on bulk RNA isolated from whole tissue; as such, we could not ascertain the cell source responsible for altered expression of *Ace2* and *Tmprss2* in the lung and trachea of obese mice. However, recent single-cell RNA sequencing analyses identified type II pneumocytes and ciliated cells as the two major cell types in the human lung that express *ACE2* and *TMPRSS2* [21–24]. (3) *Ace2* and *Tmprss2* expression levels were only assessed in ~6-month-old mice, corresponding to young human adults. COVID-19 mortality is significantly higher in people >70 years old [39]. Thus, further assessment of *Ace2* and *Tmprss2* expression in midlife (~1-year-old) and geriatric (~1.5-year-old) mice is warranted.

In summary, our study provides valuable insights into dynamic expression of the SARS-CoV-2 cell entry receptor and an important associated protease in the context of obesity and sex. This information will help inform our ongoing understanding of COVID-19 pathophysiology.

## ACKNOWLEDGEMENTS

This work was partly supported by grants from the National Institutes of Health (DK084171 to GWW).

## Abbreviations

ACE2: angiotensin converting enzyme 2
COVID-19: coronavirus disease 2019
SARS-CoV-2: severe acute respiratory syndrome coronavirus 2
TMPRSS2: transmembrane serine protease 2

## Notes

### Competing Interest Statement

The authors have declared no competing interest.

